# Individual Tree-Crown Detection in RGB Imagery Using Semi-Supervised Deep Learning Neural Networks

**DOI:** 10.1101/532952

**Authors:** Ben G. Weinstein, Sergio Marconi, Stephanie Bohlman, Alina Zare, Ethan White

## Abstract

Remote sensing can transform the speed, scale, and cost of biodiversity and forestry surveys. Data acquisition currently outpaces the ability to identify individual organisms in high resolution imagery. We outline an approach for identifying tree-crowns in RGB imagery while using a semi-supervised deep learning detection network. Individual crown delineation has been a long-standing challenge in remote sensing and available algorithms produce mixed results. We show that deep learning models can leverage existing Light Detection and Ranging (LIDAR)-based unsupervised delineation to generate trees that are used for training an initial RGB crown detection model. Despite limitations in the original unsupervised detection approach, this noisy training data may contain information from which the neural network can learn initial tree features. We then refine the initial model using a small number of higher-quality hand-annotated RGB images. We validate our proposed approach while using an open-canopy site in the National Ecological Observation Network. Our results show that a model using 434,551 self-generated trees with the addition of 2848 hand-annotated trees yields accurate predictions in natural landscapes. Using an intersection-over-union threshold of 0.5, the full model had an average tree crown recall of 0.69, with a precision of 0.61 for the visually-annotated data. The model had an average tree detection rate of 0.82 for the field collected stems. The addition of a small number of hand-annotated trees improved the performance over the initial self-supervised model. This semi-supervised deep learning approach demonstrates that remote sensing can overcome a lack of labeled training data by generating noisy data for initial training using unsupervised methods and retraining the resulting models with high quality labeled data.

## 1. Introduction

Image-based artificial intelligence can advance our understanding of individual organisms, species, and ecosystems by greatly increasing the scale and efficiency of data collection [1]. The growing availability of sub-meter airborne imagery brings opportunities for the remote sensing of biological landscapes that scales from individual organisms to global systems. However, the laborious, non-reproducible, and costly annotation of these datasets limits the use of this imagery [2].

Tree detection is a central task in forestry and ecosystem research and both commercial and scientific applications rely on delineating individual tree crowns from imagery [3,4]. While there has been considerable research in unsupervised tree detection while using airborne LIDAR (Light Detection and Ranging; a sensor that uses laser pulses to map three-dimensional structure) [3,5,6], less is known regarding tree detection in RGB (red, green, blue) orthophotos. When compared to LIDAR, two dimensional RGB orthophotos are less expensive to acquire and easier to process, but they lack direct three-dimensional information on crown shape. Effective RGB-based tree detection would unlock data at much larger scales due to increasing satellite-based RGB resolution and the growing use of uncrewed aerial vehicles. Initial studies of tree detection in RGB imagery focused on pixel-based methods and watershed algorithms to find local maxima among the pixels to create potential tree crowns [7]. When combined with hand-crafted rules on tree geometries, these approaches separately performed tree-detection and crown delineation [6,8]. The need to hand-craft tree geometry rules makes it a challenge to create a single approach that encompass a range of tree types [9].

Deep learning is a well-established method for detecting and identifying objects in RGB images, but it has only recently been applied to vegetation detection [10,11]. When compared to previous rule-based approaches, deep learning has three features that make it ideal for tree detection. First, convolutional neural networks (CNNs) directly delineate objects of interest from training data rather than using hand-crafted pixel features. This reduces the expertise that is required for each use-case and improves the transferability among projects [12]. Second, CNNs learn hierarchical combinations of image features that focus on the object-level, rather than pixel-level, representations of objects. Finally, neural networks are re-trainable to incorporate the idiosyncrasies of individual datasets. This allows for models to be refined with data from new local areas without discarding information from previous training sets.

The challenge for applying deep learning to natural systems is the need for large training datasets. A lack of training data is a pervasive problem in remote sensing due to the cost of data collection and annotation [13]. In addition, the spatial extent of training data often prohibits the field-based verification of annotated objects. For tree detection, the high variation in tree crown appearance, due to taxonomy, health status, and human management, increases the risk of overfitting when using small amounts of training data [10]. One approach to addressing the data limitation in deep learning is “self-supervised learning” (*sensus* [14]), which uses unsupervised methods to generate training data that is used to train supervised models [15]. This approach has recently been applied to remote sensing for hyperspectral image classification [9]. Self-supervision, which only relies on unlabeled data, can be combined with labeled data in a semi-supervised framework (*sensu* [14]), which may improve deep learning on limited training data by providing neural networks the opportunity to learn generalized features on a wider array of training examples, followed by retraining on a smaller number of high quality annotations [16]. It is unknown whether moderate to low quality annotations can be used to generate trees for initial model training, given the imperfect nature of existing unsupervised tree delimitation approaches.

## 2. Materials and Methods

We propose a semi-supervised pipeline for detecting tree crowns based on RGB data. Figure 1 outlines this pipeline. In the proposed workflow, a LIDAR unsupervised algorithm generates initial tree predictions. The bounding box for each tree is extracted and the corresponding RGB crop is used to train an initial deep learning model. Subsequently, using this self-supervised model as a starting point, we retrain the model using a small number of hand-annotations to correct errors from the unsupervised detection. The LIDAR data is only used to initialize the training of the network. It is not used for the final prediction step. The result is a deep learning neural network that combines unsupervised and supervised approaches to perform tree delineation in new RGB imagery without the need for co-registered LIDAR data. This provides the potential for expanding the use of deep learning in remote sensing applications with limited labeled data by exploring whether generating hundreds of thousands of noisy labels will yield improved performance, even though these labeled data are imperfect due to the limitations of the generative algorithm [17].

**Figure 1.**
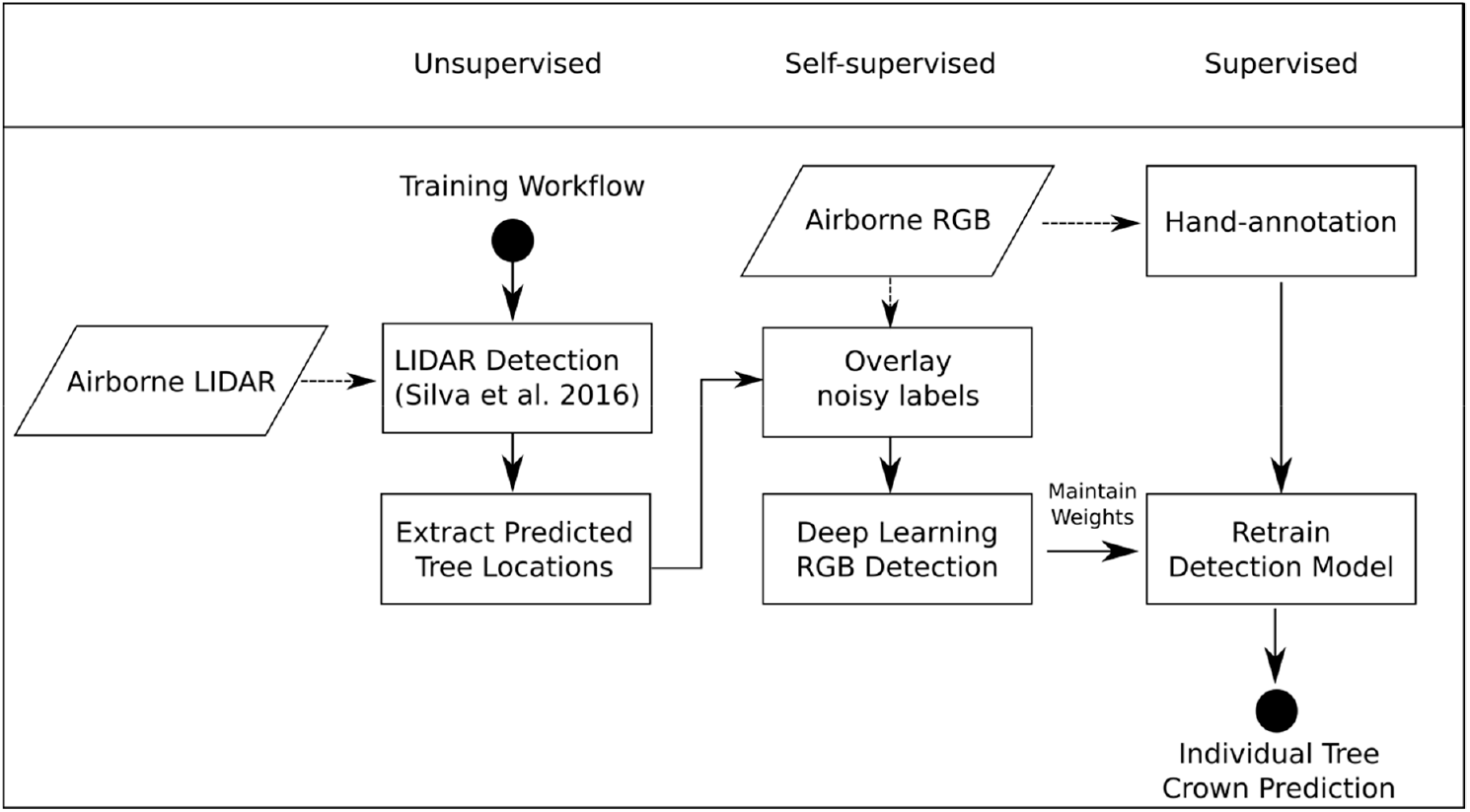
A conceptual figure of the proposed semi-supervised pipeline. A Light Detection and Ranging (LIDAR)-based unsupervised detection generates initial training data for a self-supervised (red, green, blue (RGB) deep learning model. The model is then retrained based on a small number of hand-annotated trees to create the full model.

### 2.1. Study Site and Field Data

We used data from the National Ecological Observatory Network (NEON) site at the San Joaquin Experimental Range in California to assess our proposed approach (Figure 2). The site contains an open woodland of live oak (*Quercus agrifolia*), blue oak (*Quercus douglasii*), and foothill pine (*Pinus sabiniana*) forest. The majority of the site is a single-story canopy with mixed understory of herbaceous vegetation.

**Figure 2.**
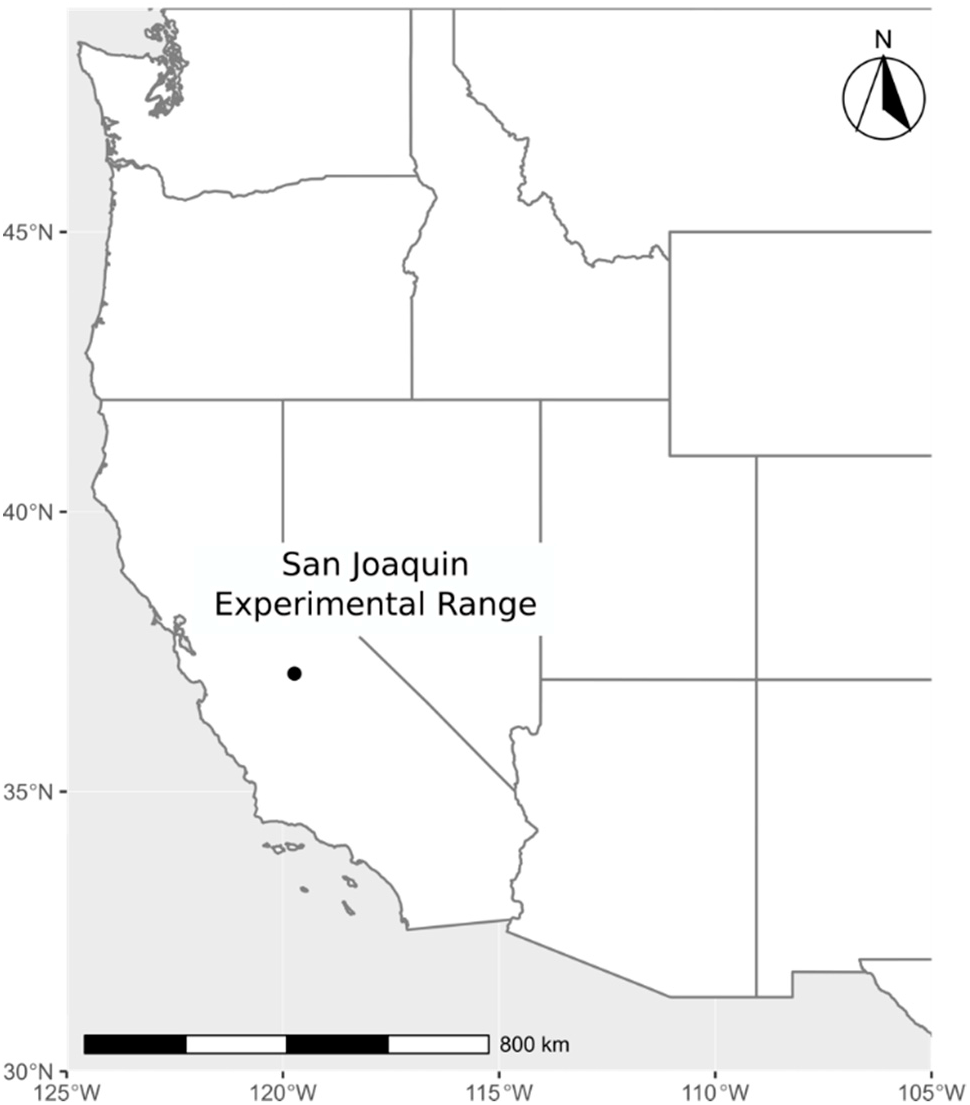
Map showing the location of the study site in the western United States.

The aerial remote sensing data products were provided by the NEON Airborne Observation Platform. We used the NEON 2018 “classified LiDAR point cloud” data product (NEON ID: DP1.30003.001) and the “orthorectified camera mosaic” (NEON ID: DP1.30010.001). The LiDAR data consist of three-dimensional (3D) spatial point coordinates (4-6 points/m^2^). which provides high resolution information regarding crown shape and height. The RGB data are a 1km × 1km mosaic of individual images with a cell size of 0.1 meters (Figure 3). Both data products are georeferenced in the UTM projection Zone 11. In addition to airborne data, NEON field teams semi-annually catalog “Woody Plant Vegetation Structure” (NEON ID: DP1.10098.001), which lists the tag and species identity of trees with diameter at breast height> 10cm in 40m × 40m plots at the site. For each tagged tree, the trunk location was obtained while using the azimuth and distance to the nearest georeferenced point within the plot. All the data are publicly available on the NEON Data Portal (http://data.neonscience.org/). All code for this project is available on GitHub (https://github.com/weecology/DeepLidar) and archived on Zenodo [18].

**Figure 3.**
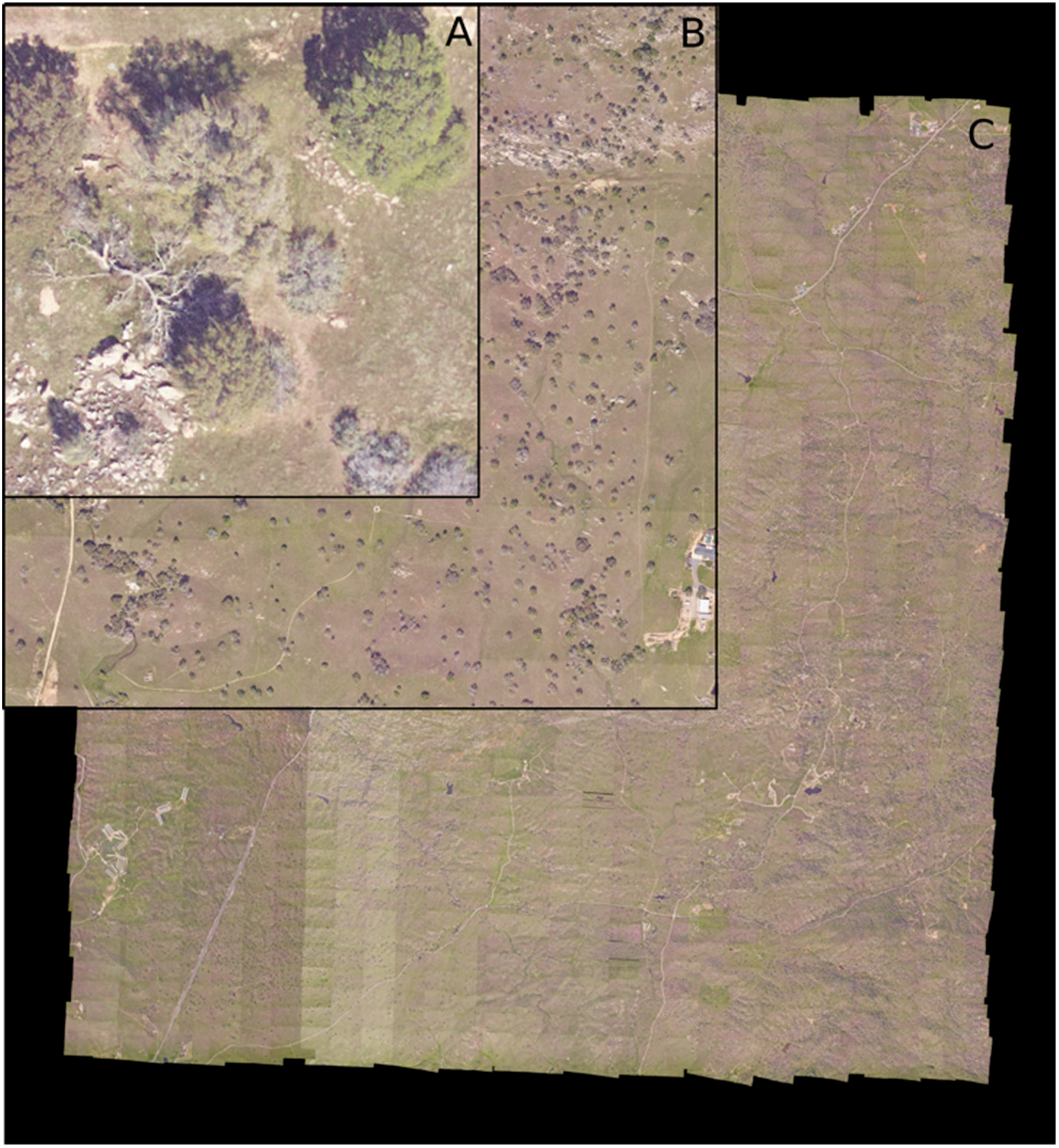
The San Joaquin, CA (SJER) site (**C**) in National Ecological Observation Network contains 148 1 km^2^ tiles (**B**), each with a spatial resolution of 0.1 m. For our analysis, we further divided each tile in 40 × 40 m windows (**A**) for individual tree prediction (n = 729 per 1 km^2^ tile).

For hand annotations, we selected two 1 km × 1 km RGB tiles and used the program RectLabel (https://rectlabel.com/) to draw bounding boxes around each visible tree. We chose not to include snags, or low bushes that appeared to be non-woody. In total, we hand-annotated 2848 trees for the San Joaquin site. In addition to the two 1 km tiles, we hand-annotated canopy bounding boxes on the cropped RGB images for each NEON field plot (n = 34), which were withheld from training and were used as a validation dataset.

### 2.2. Unsupervised LIDAR Detection

We tested three existing unsupervised algorithms for use in generating trees for the self-supervised portion of the workflow [19–21]. Existing unsupervised algorithms yield imperfect crown delineations, in part, because: 1) the algorithms are not designed to learn the specifics of different regions and datasets; 2) it is difficult to design hand-crafted features that are flexible enough to encompass the high variability in tree appearance; 3) distinguishing between trees and vertical objects, such as boulders and artificial structures, can be difficult with only three-dimensional LIDAR data. We evaluated three available unsupervised LIDAR detection algorithms while using the recall and precision statistics for hand-annotated images (see Model evaluation) in order to choose the best performing algorithm to generate training labels [19–21]. We then used the best performing method ([22]) to create initial self-supervised tree predictions in the LIDAR point cloud. This algorithm uses a canopy height model and a threshold of tree height to crown width to cluster the LIDAR cloud into individual trees (Figure 4). We used a canopy height model of 0.5m resolution to generate local tree tops and a maximum crown diameter of 60% of tree height. A bounding box was automatically drawn over the entire set of points assigned to each tree to create the training data. In total, we generated 434,551 unsupervised tree labels to use during model training. By pretraining the RGB network on these unsupervised labels, the model learns a wider variety of tree shapes and appearances than would be possible when solely using hand annotated training data.

**Figure 4.**
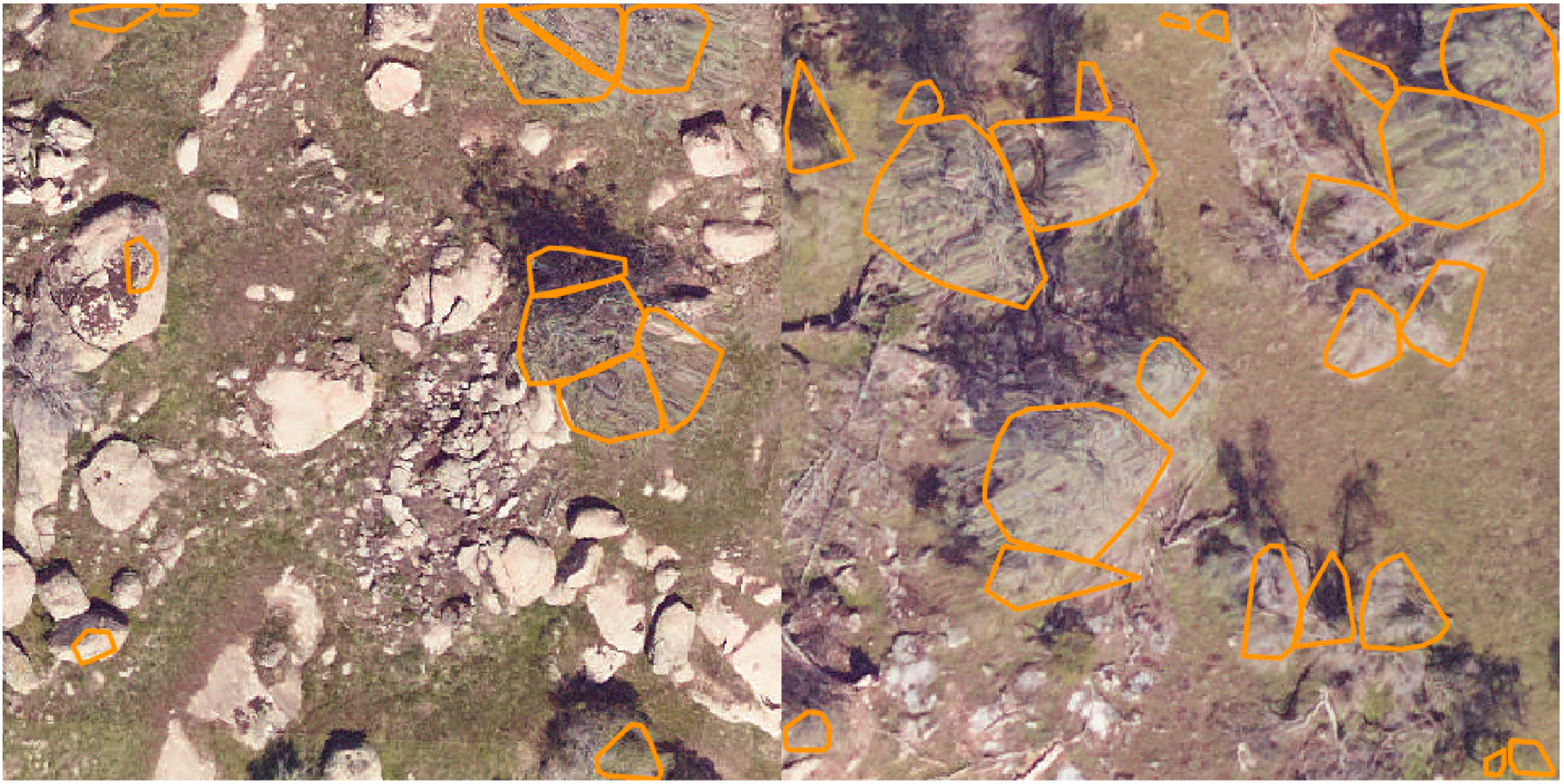
Example results from the unsupervised lidar algorithm [22], as implemented in the R liDR package [23]. Two plots from the San Joaquin National Ecological Observatory Network (NEON) site are shown (SJER_009, SJER_010).

### 2.3. Deep Learning RGB Detection

Convolutional neural networks are often used for object detection, due to their ability to represent semantic information as combinations of image features. Early applications passed a sliding window over the entire image and treated each window as a separate classification problem. This was improved by considering the potential detection boxes that are generated by image segmentation techniques [23] or by combining the bounding box proposal and classification into a single deep learning framework [24]. We chose the retinanet one-stage detector [25,26], which allows for pixel information to be shared at multiple scales, from individual pixels to groups of connected objects for learning both bounding boxes and image classes. Retinanet differs from other object detection frameworks, such as RCNN, by combining object detection and classification into a single network. This allows for faster training and it decreases the sensitivity to the number of box proposals among the images [25]. We used a resnet-50 classification backbone that was pretrained on the ImageNet dataset [27]. We experimented with deeper architectures (resnet-101 and resnet-152), but found no improvement where offset increased the training time.

We first cut the tile into smaller windows for model training, since the entire 1 km RGB tile cannot fit into GPU memory. We experimented with a number of different window sizes and found the optimal performance at 400 × 400 pixels due to a balance between memory constraints and providing the model sufficient spatial context for tree detection. We allowed each window to overlap by 5%, to ensure that we captured all trees that were potentially divided among images during cropping. This resulted in 729 windows per 1 km tile. The order of tiles and windows were randomized before training to minimize overfitting among epochs. Using the pool of unsupervised tree predictions, we trained the network with a batch size of 6 on a Tesla K80 GPU for eight epochs. For each predicted tree, the model returns a bounding box and a confidence score (0–1). After prediction, we passed each image through a non-max suppression filter to remove the predicted boxes that overlapped by more than 15%, only maintaining the box with the superior predicted score. Finally, we removed boxes within confidence scores less than 0.2.

### 2.4. Model Evaluation

We used the NEON woody vegetation data to evaluate tree recall while using field-collected points corresponding to individual tree stems (n = 111 trees). A field-collected tree point was considered to be correctly predicted if the point fell within a predicted bounding box. This is a more conservative approach than most other studies, where the field-collected tree point is considered to be correctly predicted if an edge of the bounding box falls within a horizontal search radius (e.g., 3 m in [28] to 8 m in [29]). Due to these variations in accuracy measurement, it is difficult to establish state-of-art performance, but 70–80% detection rate between the predicted trees and field located trees is typical [5,9]. There are too few previous attempts at individual tree crown prediction to provide an expectation for accuracy, given the variation in tree appearance and segmentation difficulty.

We computed recall and precision based on an intersection-over-union score of greater than 0.5 for each predicted crown to evaluate the hand-annotated crown areas. The intersection-over-union evaluation metric measures the area of overlap divided by the area of union of the ground truth bounding box and the predicted bounding box. Direct comparisons of predicted and observed crown overlap are rarely performed due to the difficulty of collecting data for a sufficient number of validation examples. The most common approach is to compare the predicted crown area to a matched tree, such as in [30] or use per pixel overlap in visually annotated data [9,31]. When compared to previous works, our use of a minimum 0.5 intersection-over-union score is more stringent. We chose this value, because it more closely resembles the required accuracy for forestry and ecological investigations [32]. Finally, to provide a baseline of comparison, we reran the evaluation data with a model trained solely using the hand-annotated data. This allows for a direct comparison of the contribution of high-quality annotations when compared to the self-supervised model or the full model combining both self-supervision and hand-annotation data.

## 3. Results

Initial exploration of existing LIDAR-based tree detection tools showed that the best performing algorithm [21] was able to correctly recall the crown area of 14% of trees at intersection-over-union score of 0.5 (Table 1). Challenges included the over-segmentation of large individual trees, erroneous predicted tree objects based on imperfections in the ground model, and the inclusion of non-tree vertical objects (Figure 5).

**Figure 5.**
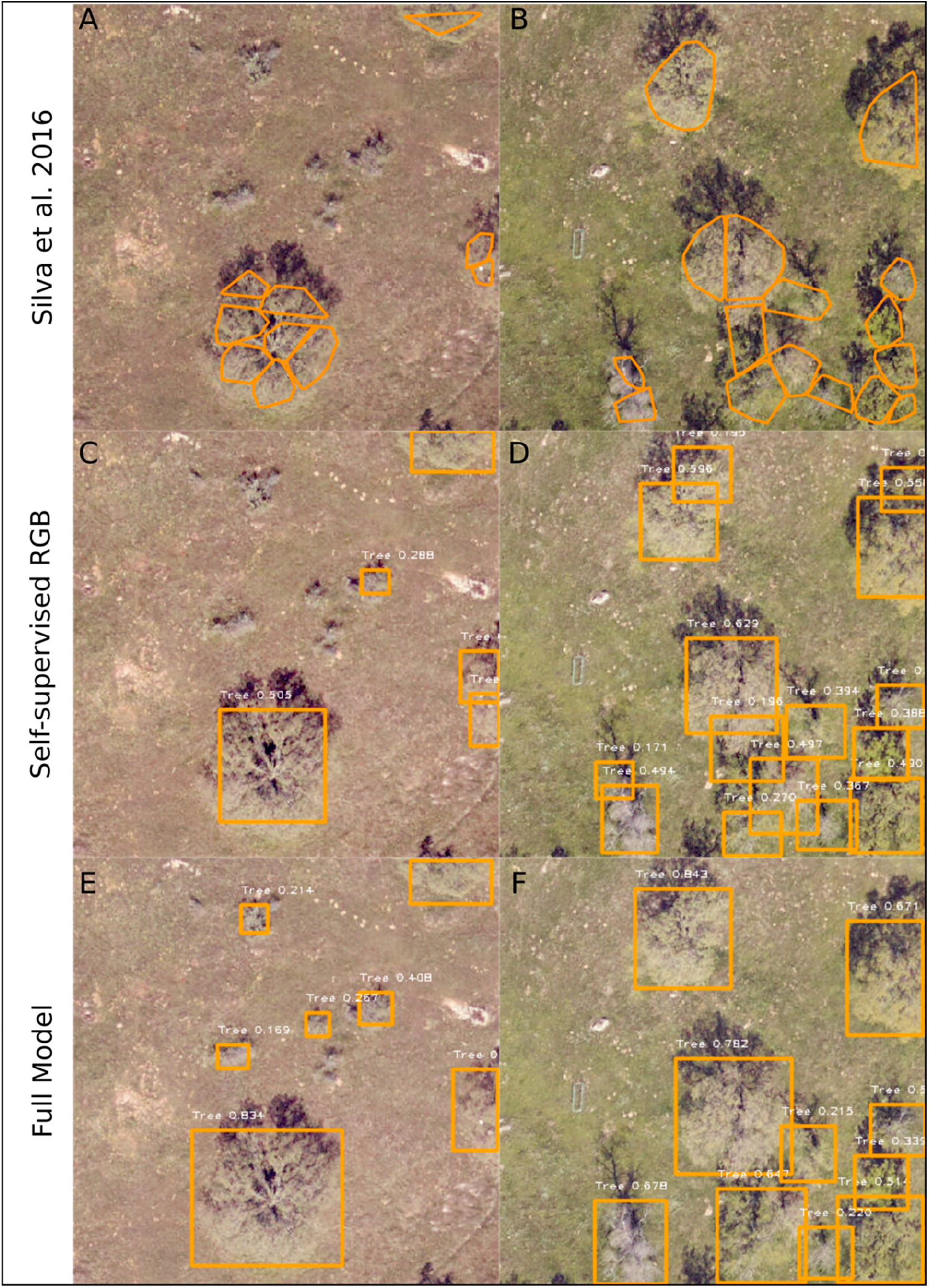
Predicted individual tree crowns for the unsupervised lidar (**A**,**B**), self-supervised RGB (**C**,**D**) and full (semi-supervised) model (**E**,**F**) for two NEON tower plots, SJER_015 (**A**,**C**,**E**), and SJER_053 (**B**,**D**,**F**) at the San Joaquin, CA site. For each tree prediction, the detection probability is shown in white.

**Table 1.** Exploratory analysis of lidar-based unsupervised algorithms. Recall and precision statistics are shown for intersection-over-union with a threshold of 0.5 overlap for the hand annotated trees on the NEON field plots (n = 271 trees).

| LIDAR Algorithm | Recall | Precision |
| --- | --- | --- |
| Li et al. (2012) [21] | 0.107 | 0.021 |
| Dalponte et al. (2016) [20] | 0.138 | 0.083 |
| Silva et al. (2016) [22] | 0.142 | 0.071 |

We extracted RGB crops and pretrained the RGB neural network using the bounding boxes from the Silva et al. (2016) predictions. This self-supervised network had a field collected stem recall of 0.83, and a hand-annotated crown area recall of 0.53 with a precision of 0.32. Retraining the self-supervised model with hand-annotated trees increased the recall of the hand annotated tree crowns to 0.69 with a precision of 0.61 (Table 2, Figure 5). The field collected stem recall did not meaningfully change among the models. We anticipate the remaining stems that were not captured are either not visible in the airborne image or belong to trees judged to be too small based on the training data.

**Table 2.** Evaluation metrics for each of the models. All evaluation was conducted on the 34 NEON field plots. Stem recall was calculated using the field-collected tree stem locations (n = 111 trees). Precision and recall for crown overlap was calculated on hand-annotated bounding boxes around each tree crown (n = 271 trees) with a minimum predicted probability threshold of 0.5.

| Model | Hand-annotated crown overlap (>50%) |  | Stem Recall |
| --- | --- | --- | --- |
|  | Recall | Precision |  |
| Silva et al. 2016 | 0.14 | 0.07 | 0.79 |
| Hand-annotation only | 0.38 | 0.60 | 0.79 |
| Self-supervised RGB | 0.53 | 0.32 | 0.83 |
| Full Model | 0.69 | 0.61 | 0.81 |

By comparing the images of the predictions from the unsupervised LIDAR detection, the self-supervised RGB deep learning model, and the combined full model, we can learn about the contributions of each stage of the pipeline. The LIDAR unsupervised detection does a good job of identifying trees versus background based on height. Most of the small trees are well segmented, but there is consistent over-segmentation of the large trees, with multiple crown predictions abutting together. Visual inspection shows that these predictions represent multiple major branches of a single large tree, rather than multiple small trees (Figure 5A). These large trees are more accurately segmented in the self-supervised RGB model, but there is a proliferation of bounding boxes, and overall lower confidence scores, even for well-resolved trees (Figure 5D). This is shown in the precision-recall curves for the hand-annotated validation data, which has a higher level of precision for the same level of bounding box recall (Figure 6).

**Figure 6.**
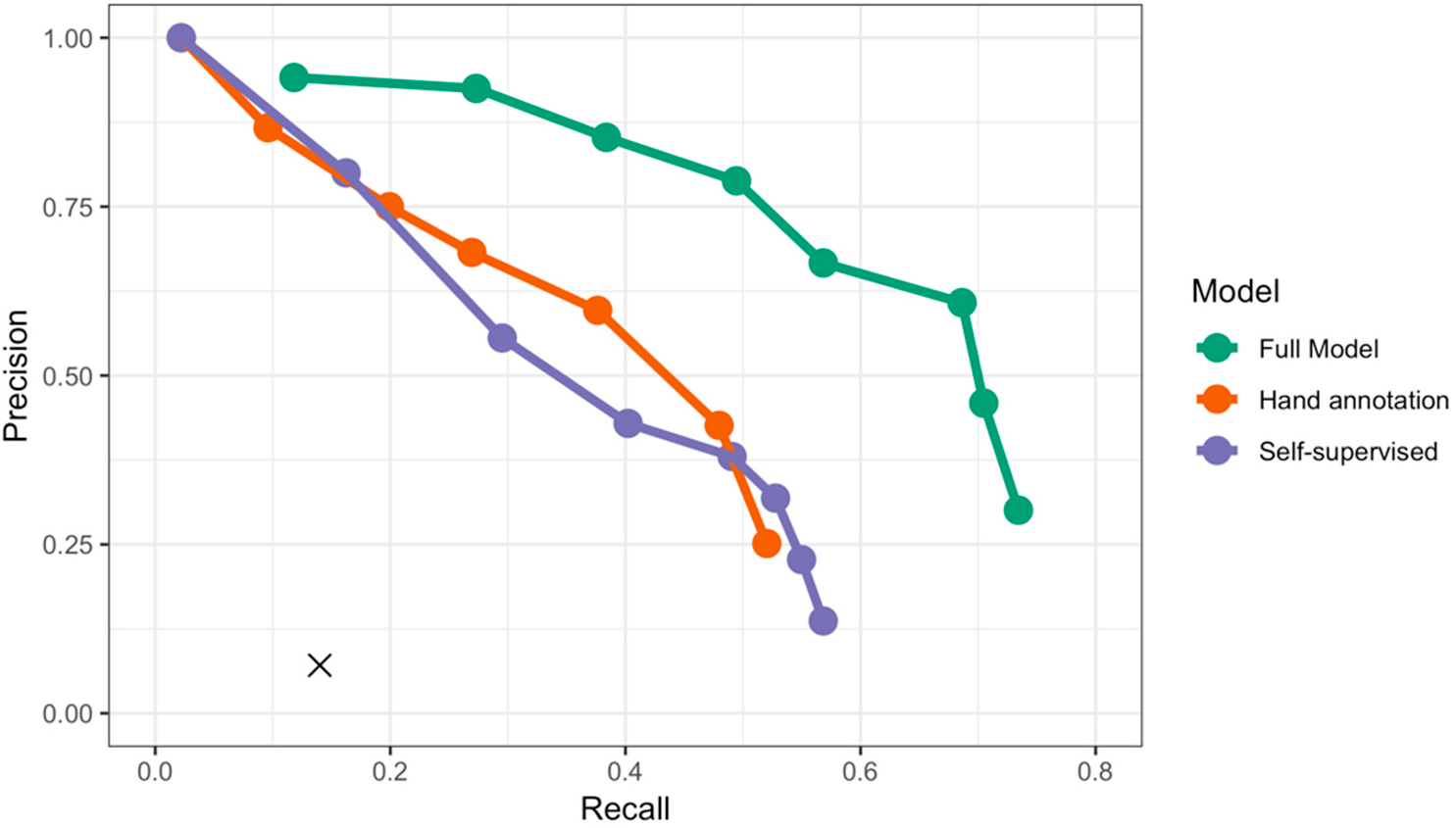
Precision-recall curves for the hand-annotated NEON plots. For each model, we calculated the proportion of correctly predicted boxes for score thresholds [0,0.1, …,0.7]. An annotation was considered to be correctly predicted if the intersection-over-union (IoU) score was greater than 0.5. The recall and precision scores for the initial lidar-based unsupervised algorithm is shown in black X.

The full model reduces the extraneous boxes and improves the segmentation of large trees by combining the self-supervised and the hand annotated datasets (Figure 7). The full model has optimal performance in areas of well-spaced large trees (Figure 7B), but it tends to under-segment small clusters of trees (Figure 7C).

**Figure 7.**
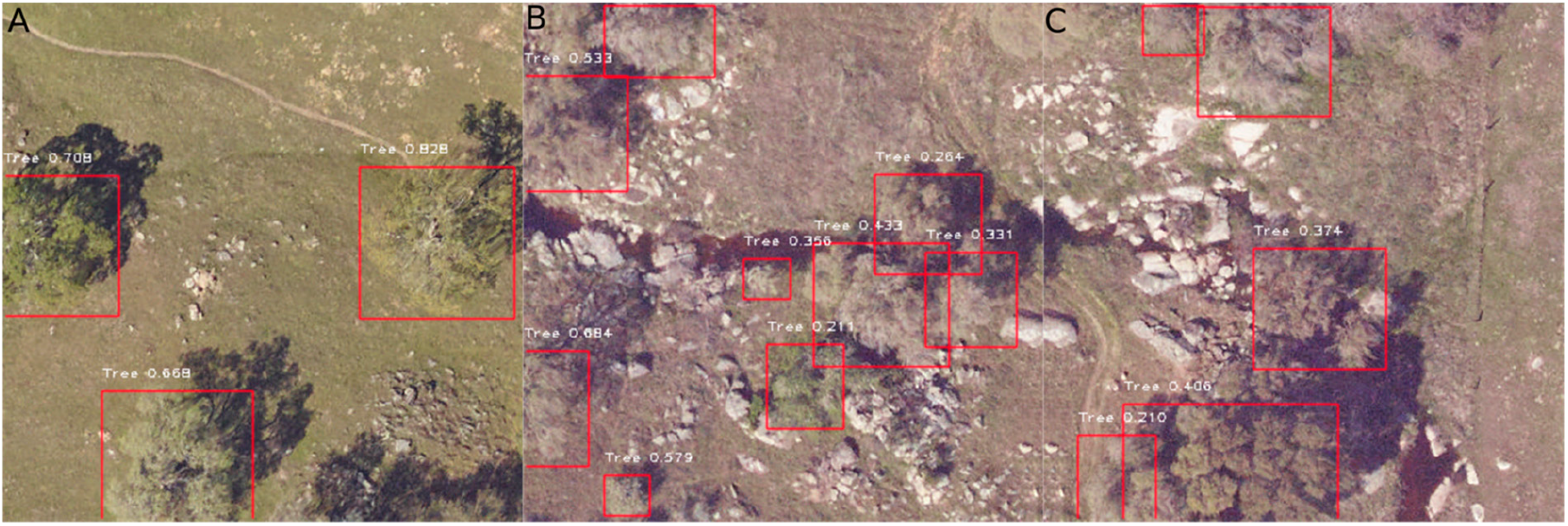
Predictions from the full model on data withheld from model training. Canopy complexity increases from (**A**) well-defined large trees to (**B**) mixed-species canopies to (**C**) tightly packed clusters of trees. As canopy complexity increases, the full model tends to under-segment small tree clusters.

## 4. Discussion

We built a neural network-based pipeline for identifying individual trees in RGB imagery using recent developments in deep learning. Commercial high resolution RGB data is increasingly available at near global scales, which means that accurate RGB based crown delineation methods could be used to detect overstory trees at unprecedented extents. We used an unsupervised LIDAR tree detection algorithm to generate labels for initial training to address the long-standing challenge of a lack of labeled training data. This self-supervised approach allows for the network to learn the general features of trees, even if the LIDAR-based unsupervised detection is imperfect. The addition of only 2848 hand-annotated trees generated a final model that performed well when applied to a large geographic area. This approach opens the door for the use of deep learning in airborne biodiversity surveys, despite the persistent lack of annotated data in the forestry and ecology datasets.

Many of the false positives in our evaluation dataset were due to disagreements between the hand annotations, unsupervised LIDAR pretraining, and RGB prediction in what defines a tree. For example, small trees were often considered too low for inclusion in the LIDAR algorithm (Figure 5A), whereas they were included in the full model based on the hand-annotations (Figure 5B). Similarly, large bushes were sometimes included in hand annotations due to the difficulty of determining the overall woody structure. When deploying these models to the applied problems, it will be important to have strict quantitative guidelines that define tree definitions. Where LIDAR data is available, draping the two-dimensional (2D) boxes over the 3D point cloud to filter out the points based on vertical height should be useful for improving precision. It should be noted that the quantitative results are likely biased toward the RGB model, since looking at the RGB, and not the LIDAR data, made the hand-annotations. However, the good recall rate for the field-collected stems suggests that hand annotations were useful in capturing field conditions. An unexpected benefit of the RGB model was the ability to discriminate trees from other vertical objects, such as houses or poles, despite a lack of distinction in the unsupervised LIDAR training data (Figure 8). This may be useful in urban tree detection and other non-forested sites.

**Figure 8.**
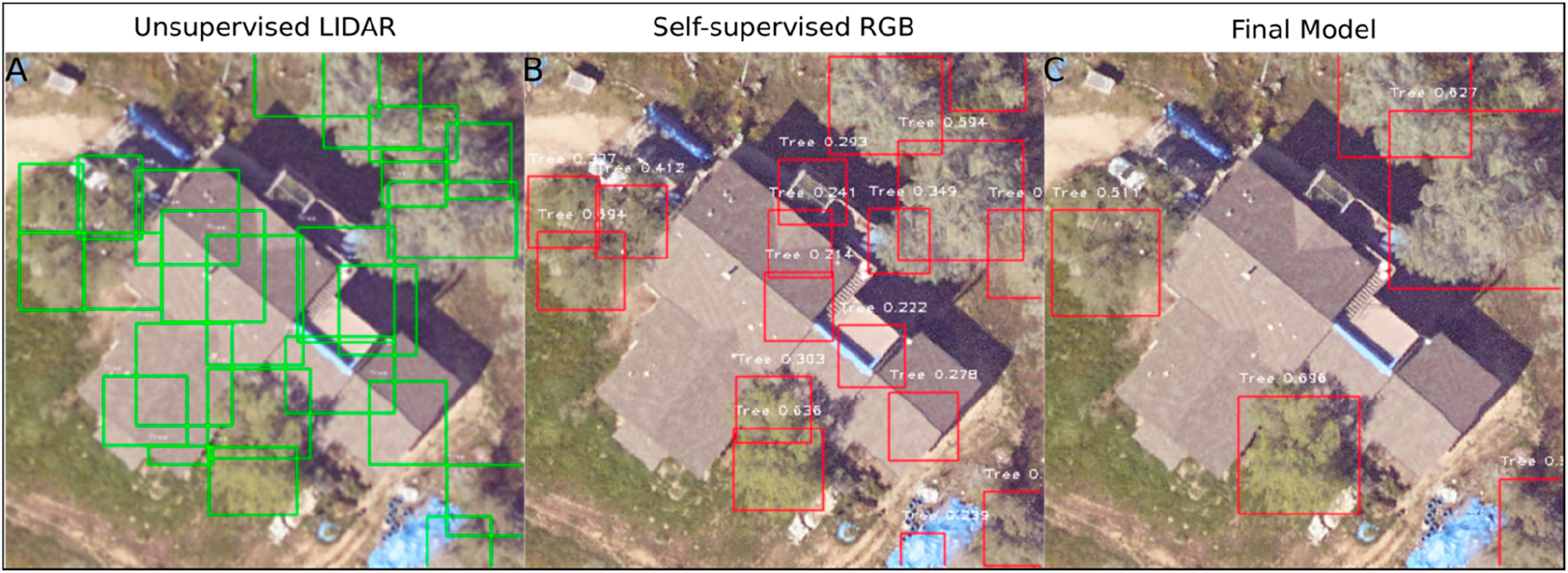
Improvement in prediction quality during the training pipeline. (**A**) Bounding boxes from the lidar-based unsupervised detection erroneously identified artificial structures as trees. (**B**) Predictions from the self-supervised RGB model showed that the addition of RGB data diminished the effect of incorrectly labeled training data, with only edges of the artificial structures maintained as tree predictions. (**C**) In the full semi-supervised model, combining the self-supervised RGB data with hand-annotations eliminated the influence of the original misclassification in the training data, while still capturing the majority of trees in the image.

It is likely that accurate tree detection will be region specific, and that the best model will vary among environments. This will require training a new model for each geographic area while using both RGB and LIDAR training data. The proposed approach could save resources by allowing a smaller scale LIDAR flight to generate training data, and then cover a much larger area with less expensive RGB orthophotos. Uncrewed aerial vehicles (UAVs) can be used for capturing LIDAR at high resolution, but at a limited spatial extent. In combination with our method, these UAVs may allow for the cost effective development of custom regional tree detection models. In addition, the National Ecological Observatory Network, which provided the data for this analysis, has 45 forested NEON sites that were selected to cover the major ecoclimatic domains in the United States. These sites could serve as pools of LIDAR and RGB data at 10,000 ha scales for regional model training. Combining these two detectors together could produce accurate individual level tree maps at broad scales, with potential applications in forest inventory, ecosystem health, post-natural disaster recovery, and carbon dynamics.

While the semi-supervised deep learning method performed well at the open-canopy test site, geographic areas with complex canopy conditions will be more challenging. As the scale and tree diversity increase, the model will need additional hand-annotated training data to capture the variation within the tree class. We found that approximately 2000 trees produced reasonable performance in the oak woodland landscape while there is no hard rule for the minimum hand-annotations needed. It is difficult to diagnose the exact nature of the features used for tree detections, given the complex nature of convolutional neural networks. One potential is that the model detects shadows of vertical objects and uses them to help locate crown areas. Whether these features generalize to more complex canopy structures and new ecosystems remains an open question.

The current model only uses LIDAR in the pretraining step. Where available, directly incorporating a LIDAR canopy height model into the deep learning approach should allow the model to simultaneously learn the vertical features of individual trees, in addition to the two-dimensional color features in the RGB data. Recent applications of three-dimensional CNNs [33], as well as point-based semantic segmentation [34], provide new avenues for joint multi-sensor modeling. These developments will be crucial in segmenting the complex canopies that overlap in the two-dimensional RGB imagery. In addition, recent extensions of region-proposal networks refine the bounding boxes to identify the individual pixels that belong to a class [35]. This will provide a better estimate of the tree crown area, as trees typically have a non-rectangular shape.

## 5. Conclusions

Applying deep learning models to natural landscapes opens new opportunities in ecology, forestry, and land management. Despite a lack of high-quality training data, deep learning algorithms can be deployed for tree prediction while using unsupervised detection to produce generated trees for pretraining the neural network. Although the lidar-based algorithm that was used to generate the pretraining data achieved less than 20% recall of hand-annotated tree crowns, the deeply learned RGB features from those data achieved greater than 50% recall. When combined with a small number of hand-annotated images, the recall increased to 69% with 60% precision. As shown by the comparison with field-collected stems, the majority of the remaining predictions represent valid trees (>80%), but the overlap with the hand-estimated crown area was less than the desired 50%. Many previous papers have used a lower overlap threshold (e.g., 20% overlap in [36]), and we expect this value to improve with a combination of better validation data and more hand-annotated training samples.

There is the potential for this method to provide additional important information regarding natural systems, in addition to scaling tree detection at much lower costs. The current model could be expanded from a single class, “Tree”, to one that provides more detailed classifications that are based on taxonomy and health status. For example, splitting the “Tree” class into living and dead trees would provide management insight when surveying for outbreaks of tree pests and pathogens [37], as well as post-fire timber operations [38]. The growing use of drones in environmental remote sensing also opens up additional possibilities of combining high density local information with broad scale information captured from traditional fixed wing and satellite mounted sensors. Unsupervised tree detection algorithms have been shown to be more effective at very high point densities (e.g., [36]). We anticipate that, as the quality of the unsupervised algorithm data increases, fewer hand annotated samples will be needed to customize the model to a local geographic area. In addition, the availability of hyperspectral data could assist in dividing the ‘tree’ class into multiple species labels. This would yield additional insights into the economic value, ecological habitat, and carbon storage capacity for large geographic areas [39]. While further work is needed to understand the best way to combine data among scales and sensors, we show that deep learning-based approaches hold the potential for large scale actionable information on natural systems to be derived from remote sensing data.

## Author Contributions

B.G.W., E.W., S.B. and A.Z. conceived of project design. E.W. and S.M. collected the preliminary data. B.G.W. performed the analysis and wrote the text. All authors contributed to the text.

## Acknowledgments

Financial support for S.A.B was provided by the USDA/NIFA McIntire-Stennis program (FLA-FOR-005470).and a sabbatical fellowship award from sDiv (the Synthesis Centre of iDiv; DFG FZT 118).

